# Reward timing and its expression by inhibitory interneurons in the mouse primary visual cortex

**DOI:** 10.1101/785824

**Authors:** Kevin J Monk, Simon Allard, Marshall G Hussain Shuler

**Affiliations:** The Solomon H. Snyder Department of Neuroscience, The Johns Hopkins University School of Medicine, 725 N Wolfe Street, Baltimore, MD 21205; Kavli Neuroscience Discovery Institute, The Johns Hopkins University School of Medicine, 725 N Wolfe Street, Baltimore, MD 21205

## Abstract

Primary sensory cortex has historically been studied as a low-level feature detector, but has more recently been implicated in many higher-level cognitive functions. For instance, after an animal learns that a light predicts water at a fixed delay, neurons in primary visual cortex (V1) can produce “reward timing activity” (i.e., spike modulation of various forms that relate the interval between the visual stimulus and expected reward). The manner by which V1 produces these representations is unknown. Here, we combine behavior, *in vivo* electrophysiology, and optogenetics to investigate the characteristics of and circuit mechanisms underlying V1 reward timing in the head-fixed mouse. We find that reward timing activity is present in mouse V1, that inhibitory interneurons participate in reward timing, and that these representations are consistent with a theorized network architecture. Together, these results deepen our understanding of V1 reward timing and the manner by which it is produced.

## Introduction

Primary sensory cortex is classically regarded as a low-level feature detector providing simple representations for higher-order areas. In the visual system, representations in early areas relate to simple features, and through the cortical hierarchy, these signals are transformed into complex representations of the external world (Hubel and Wiesel, 1959, 1965; Felleman and Van Essen, 1991). Yet, it is becoming increasingly clear that representations in primary sensory areas are updated when stimuli gain meaning through associative learning (McGann, 2015). Specifically, this is seen in primary gustatory (Vincis and Fontanini, 2016), somatosensory (Wiest et al., 2010; Gdalyahu et al., 2012; Pais-Vieira et al., 2013), auditory (Polley et al., 2004; Rutkowski and Weinberger, 2005; Guo et al., 2019), and visual (Shuler and Bear, 2006; Gavornik and Bear, 2014a, 2014b; Makino and Komiyama, 2015; Goard et al., 2016; Goltstein et al., 2018) cortices, and in the olfactory bulb (Kay and Laurent, 1999; Kass et al., 2013; Ross and Fletcher, 2018).

Previous work shows that individual neurons in rodent primary visual cortex (V1) express reward timing activity (Shuler and Bear, 2006; Chubykin et al., 2013; Liu et al., 2015; Zold and Hussain Shuler, 2015). Reward timing activity is a representation of time between a visual stimulus and a reward, expressed as one of three forms: 1) a sustained increase (SI) or 2) sustained decrease (SD) of activity until the time of expected reward, or 3) a peak (PK) of activity around the time of the expected reward. Prior studies advance V1 as a substrate in the learning of this timing activity and implicate acetylcholine as a reinforcing signal: lesions of cholinergic axons within V1 block the ability for V1 to learn reward timing activity (Chubykin et al., 2013); pairing visual stimuli with local activation of cholinergic fibers in V1 mimics behaviorally-conditioned reward timing (Liu et al., 2015); V1 timing activity correlates with timing behavior and perturbation of this activity lawfully shifts timing behavior (Namboodiri et al., 2015; Levy et al., 2017).

These observations implicate V1 as a substrate for learning reward timing activity—thought to occur through a reinforcement learning process (Hussain Shuler, 2016)—but it remains unclear how V1 circuitry produces reward timing activity with its various response forms. A computational model proposes a solution with a specific connectivity motif within a network of excitatory and inhibitory cells (Huertas et al., 2015). This network architecture has two implications: (1) inhibitory interneurons represent time predominantly as the SI form and (2) neurons inhibited by interneurons represent time predominantly as the SD or PK form. In this network, interneurons are treated as one, monolithic group. However, V1 interneurons fall mainly into one of three subpopulations expressing either parvalbumin (PV), somatostatin (SOM), or vasoactive intestinal polypeptide (VIP) (Xu et al., 2010; Tremblay et al., 2016). Each are unique in their connectivity patterns (Pfeffer et al., 2013) and are functionally distinct during stimulus representation in V1 (Atallah et al., 2012; Lee et al., 2012; Wilson et al., 2012). It is unknown if either of the model’s implications are borne out *in vivo* and, if so, how the diversity of interneuron subtypes intersects with these implications.

Here we enrich our understanding of V1 reward timing by investigating the manner by which mouse V1 neurons produce reward timing activity, how different interneuron populations express and aid in the production of this activity, and how well biological and computational data accord with one another. In doing so, we find that V1 neurons express reward timing in a manner consistent with a theorized network architecture and that PV+ interneurons fulfill the expectations of the theorized inhibitory population.

## Results

The means by which the primary visual cortex produces the various forms of reward timing observed is unknown. We have investigated potential mechanisms through an in-depth characterization of reward timing in the mouse primary visual cortex and how this reward timing activity is expressed by inhibitory interneurons (Figure 1A for a recording schematic). Specifically, we use mice which selectively express channelrhodopsin-2 (ChR2) in interneuron subpopulations to determine how these different cell types produce and aid in the production of reward timing activity (Figure 1B). Finally, with these data we sought to compare the activity of interneuron populations with a proposed network architecture which replicates reward timing activity (Figure 1C, (Huertas et al., 2015)).

**Figure 1:**
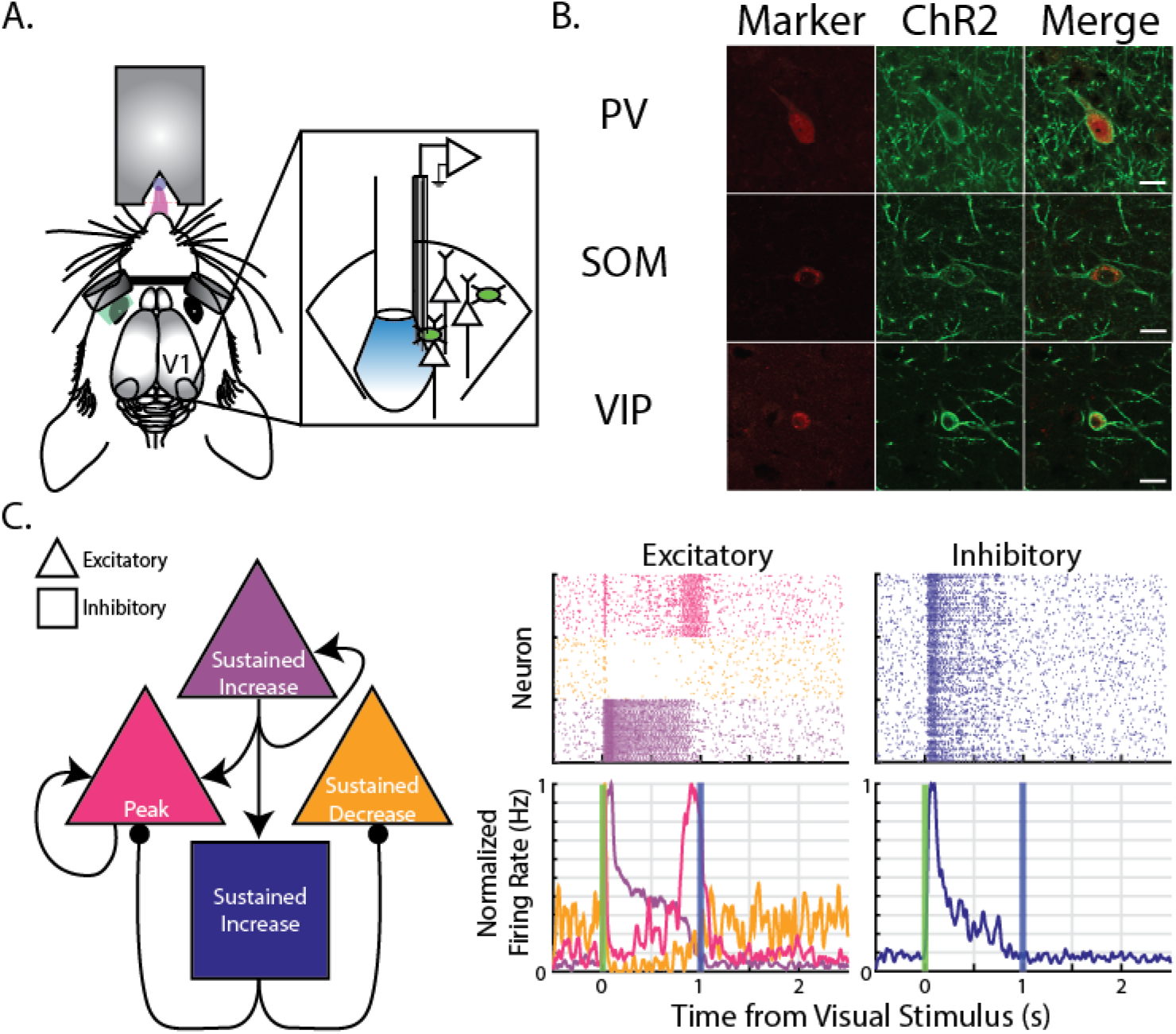
Summary of recording strategy and motivation to investigate inhibitory interneurons. (A) Cartoon schematic showing recording design. Head-fixed mice were required to lick following presentation of visual stimuli to receive a water reward after a fixed delay. Simultaneously, electrophysiological recordings were made in infragranular primary visual cortex. Depending on genotype, mice expressed channelrhodopsin-2 in inhibitory interneurons (schematized in green). (B) Channelrhodopsin-2 (ChR2) expression co-localizes with expected interneuron marker given animal’s genotype. Specifically, in PV-ChR2 animals, there was co-expression of PV and ChR2 (top row), in SOM-ChR2 animals there was co-expression of SOM and ChR2 (middle row), and in VIP-ChR2 animals, there was co-expression of VIP and ChR2 (bottom row). Scale bars in right column represent 151Jm. (C) <u>Left:</u> Schematic of the theorized network architecture. This architecture is composed of three populations of excitatory cells (triangles) and one population of inhibitory cells (square). Right: Following training of the model, simulated excitatory neurons produce reward timing responses in one of three forms (left), and simulated inhibitory units produce Sustained Increase reward timing responses. Schematic on left adapted from Huertas et al., 2015; plots on right simulated as in Huertas et al., 2015.

### Head-Fixed Mice Associate a Visual Stimulus with a Delayed Reward

Prior to investigating the circuit mechanisms by which primary visual cortex (V1) produces reward timing, we first established its existence within the head-fixed mouse preparation. Mice were trained to associate a water reward with visual stimuli (see Methods). Briefly, head-restrained mice received a 100ms visual stimulus delivered to the left or right eye with equal probability (Cue 1 and Cue 2, respectively) via head-mounted goggles and received water from a lick port placed within reach of the tongue. Trials were initiated after a lapse of time comprising a randomly selected interval and a second random interval less than the ITI during which the animal must not lick (a “lick lockout” interval). If an animal licked during this lick lockout, the lockout timer would restart. Such an ITI encourages mice to cease licking and to wait for the onset of the next trial. Upon the initiation of a trial, animals received a monocular visual stimulus delivered to the left or right eye with equal probability, after which the animal was required to make at least one lick within the subsequent delay period so that reward could be delivered at the end of the delay. On half of these trials, if the animals met this behavioral requirement, they received a small water reward (∼2µL) at the end of the conditioned interval (so-called “paired” trials). On the other half of these trials, regardless of lick behavior, reward was withheld (“catch” trials). On 20% of trials, neither a visual stimulus nor a reward were delivered although the intertrial interval and lick lockout periods expired successfully; these trials are referred to as “sham” trials and are used to verify that animals are using visual stimuli to guide licking behavior (as opposed to timing lick bouts from events other than a visual cue). The delay time used was the same for both visual stimuli within a recording session and varied across days, as follows: on short delay sessions the delay time was 1 second following visual stimulus offset, and on long delay sessions the delay time was 1.5 seconds following visual stimulus offset. A task schematic is shown in Figure 2A and behavior from an example session is shown in Figure 2B. Regardless of trial type, trials in which the animal made a lick during the delay window are defined as Hit trials and trials in which the animal did not lick were referred to as Miss trials. All data presented here, unless otherwise noted, are from Catch+Hit trials (i.e., trials in which the animal received a visual stimulus, licked during the delay, and did not receive a reward).

**Figure 2:**
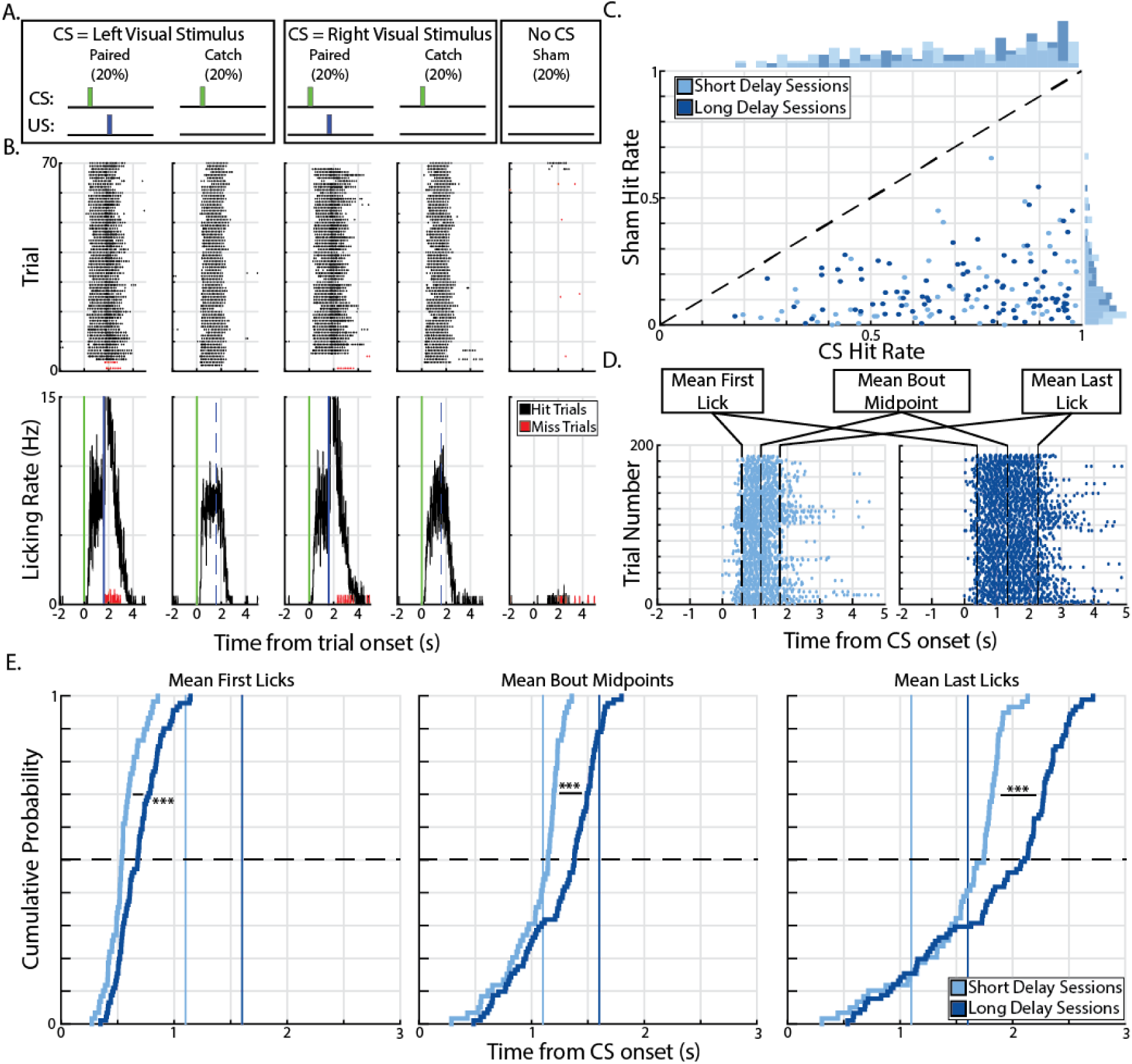
Head-fixed mice learn the reward timing task. (A) Animals were trained that a monocular visual stimulus (the conditioned stimulus,CS) delivered to the left or right eye predicted a water reward (the unconditioned stimulus,US) at a fixed delay. Upon receipt of the CS, animals were required to lick during the delay period so that reward could be delivered at the end of the short or long delay. Trials in which the reward is delivered are Paired trials and trials in which the reward iswithheld are Catch trials. Trials in which no CS nor US were delivered are Sham trials and are used to determine the ability of the animal to use the visual stimulus to guide licking behavior. (B) Example licking behavior for a single animal during a single session of the task for all trialtypes with respect to the onset of the CS (time 0, green line). Blue lines represent time of reward delivery (solid line representing the receipt of reward on paired trials and dashed line representing expected time of reward on catch trials). Top: raster plot of licking behavior where black dots are licks recorded during “Hit” trials (i.e., trials in which the animal licked during the delay window) and red dots are licks recorded during “Miss” trials (i.e., trials in which the animal did not lick in the delay window). Bottom: PSTH’s calculated from licking activity, color scheme as noted above. (C) Scatter plot of session hit rates (probability of licking on a given trial) for all trials in which the CS is delivered (CS Hit Rate) and for all trials inwhich the CS was withheld (Sham Hit Rate) for short delay sessions (light blue) and long delay sessions (dark blue). Histograms of hit rates for each trial category shown in margins and unity line is the black, dashed line. (D) Behavioral measurements (labelled,black dashed lines) for example licking behavior during Catch+Hit trials from a short delay session (left, light blue raster) and a long delay session (right, dark blue raster). (E) Cumulative distribution plots for the three behavioral measurements measured during Catch+Hit trials during short delay sessions (light blue) and long delay sessions (dark blue). Vertical lines represent reward time and horizontal line indicates the median values.*** - p < 0.001,Wilcoxon rank-sum test.

As expected, animals showed a high probability of licking in the delay period on trials where a visual stimulus was delivered (“CS trials”) and a low probability of licking during the sham trials (70.67% and 14.04%, respectively, Figure 2B and 2C). There is a significant effect of trial type (i.e., CS trial vs sham trial, χ^2^(1, 286) = 464.11, p = 7.83 × 10^−62^, Kruskal-Wallis test) on the probability that an animal licks while there is neither a significant effect of session number nor a significant interaction (Session Number: χ^2^(8, 286) = 0.51, p = 0.85; Interaction: χ^2^ (8, 286) = 1.26, p = 0.26, Kruskal-Wallis test). These results demonstrate that animals lick in response to reward-predicting visual stimulation and that their behavior had reached asymptotic performance at the time of recording.

We next addressed whether the animals time their behavioral response. To quantify the timing of the licking behavior, we made three measurements: the time of the first lick in a bout, the time of the last lick in a bout, and the mean time between the first and last lick in a bout (Bout Midpoint). The Bout Midpoint is derived from the initiation and cessation of licking, and so is not an independent measure. Rather, its inclusion is simply to determine whether the centering of lick bouts is in good accordance with the expected time of reward. These values were recorded across trials and an average of these values were calculated for a given trial type on a given day (Figure 2D for example sessions). When we compare these values, we find that the lick initiation and cessation times (and, consequently, the Bout Midpoint) are significantly smaller for short delay sessions compared to long delay sessions (Figure 2E, Mean First Licks: Z = −6.09, p = 1.11 × 10^−9^; Mean Bout Midpoints: Z = −6.71, p = 2.01 × 10^−11^; Mean Last Licks: Z = −5.73, p = 9.89 × 10^−9^, Wilcoxon rank-sum test) indicating that animals adapt their licking behavior based on the expected time to reward.

### Neurons in Primary Visual Cortex of the Head-Fixed Mouse Express Reward Timing Activity

These behavioral data indicate that animals express an internal sense of the time interval between the visual stimulus and the water reward. To determine what, if any, neural representation of time was present in V1, we recorded single unit activity bilaterally during behavioral sessions. Previous work from our lab has shown that, in similar tasks, neurons in V1 of freely-moving rats and mice represent the time interval to an expected reward in one of three forms: a sustained increase (SI) or sustained decrease (SD) of activity until the time of reward, or as a peak (PK) of activity around the time of reward (Shuler and Bear, 2006; Chubykin et al., 2013; Liu et al., 2015). These response forms were also observed here in the head-fixed mouse (Figure 3A). Using these response forms, we manually classified in a blinded fashion the peristimulus time histograms (PSTHs) of neurons for both Cue 1 and Cue 2 Catch+Hit trials (i.e., trials in which the animal received a visual stimulus, licked during the delay window, and did not receive a water reward at the end of that delay; see Methods). PSTHs created from Sham+Hit trials (that is, trials in which neither CS nor US was delivered, but had licks within the delay window) were also blindly classified as a control. Neurons could be classified as responsive during any of these trial types; as such, we began our analyses by quantifying “neural records” (i.e., a given pattern of activity a neuron produced during a trial type).

**Figure 3:**
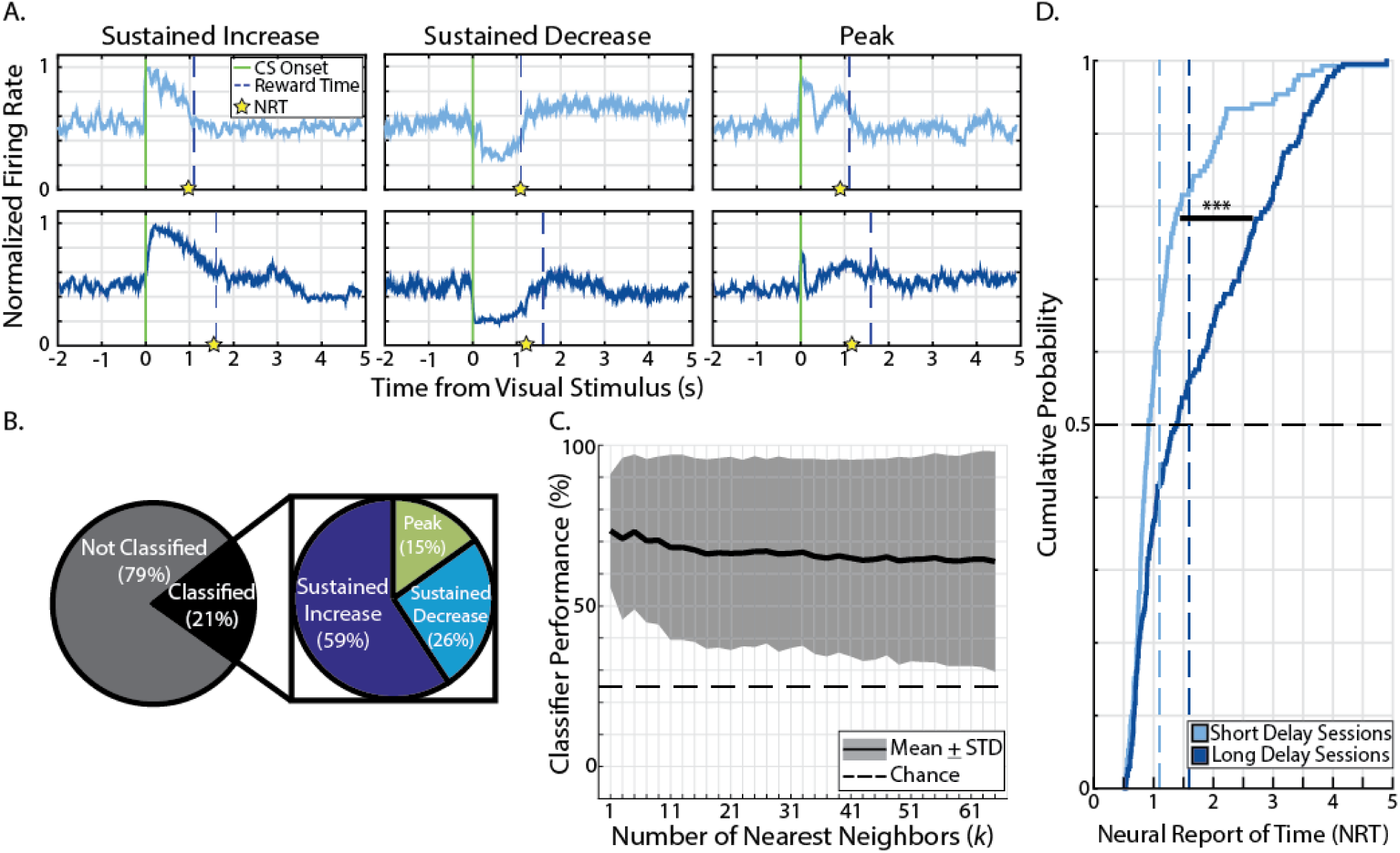
Neurons in primary visual cortex express reward timing activity in three forms. (A) Six example neurons representing the three forms of reward timing: Sustained Increase (left), Sustained Decrease (middle), and Peak (right) recorded from short delay sessions (light blue, top row) or long delay sessions (dark blue, bottom row). Activity is recorded from Catch+Hit Trials, normalized by AUC, and is plotted with respect to CS onset (green, vertical line). Dashed, vertical lines represent the time of expected reward (i.e., the conditioned interval for the session). Calculated neural report of times (NRT) for these example neurons are shown as yellow stars on x-axis. (B) Pie charts showing proportion of responses which express reward timing (left) and, of those classified responses, the proportion of the three forms of reward timing (right). (C) Performance of kNN classifier across a range of *k* nearest neighbors. Thick black line represents average performance of classifier, shaded region represents the mean± standard deviation. Dashed horizontal line represents chance performance. (D) Cumulative distribution plots of the calculated neural reports of time calculated from Catch+Hit trials during short delay sessions (light blue) and long delay sessions (dark blue). Vertical lines show time of reward and horizontal line shows median values for distributions. *** -p < 0.001, Wilcoxon rank-sum test.

We recorded from 996 neurons in the primary visual cortex which yielded 1,992 neural records from Catch+Hit trials (each neuron produced two neural records: one in response to Cue 1 and one in response to Cue 2). Of these 1,992 neural records from Catch+Hit trials, 410 (20.58%) were classified as expressing reward timing (i.e., were classified as SI, SD, or PK). These 410 records were expressed by 253 neurons (25.40% of the total recorded population). Among the 410 records: 243 (59.27%) were classified as SI (47 neurons classified for one CS, 98 neurons classified for both CS’s), 105 (25.61%) were classified as SD (33 neurons classified for one CS, 36 neurons classified for both CS’s), and 62 (15.12%) were classified as PK (30 neurons classified for one CS, 16 neurons classified for both CS’s). Only 11 of 996 (1.10%) of the neural records from Sham+Hit trials were classified as one of the forms described above. Figure 3B shows the proportions of neural records classified. We cross-validated these classifications using a *k*-Nearest Neighbors (kNN) classifier that was trained on data from even trials to predict the reward timing expression of data from odd trials (see Methods). The classifier predictions matched well with the manual classification across a range of parameters (1 ≤ *k* ≤ 65, Figure 3C) arguing that neural activity within mouse V1 fall into distinct classes of reward timing activity, as previously reported (Shuler and Bear, 2006; Chubykin et al., 2013; Liu et al., 2015).

Previous reward timing studies have shown that V1 reward timing corresponds to the delay to reward (Shuler and Bear, 2006; Chubykin et al., 2013). By ascribing to each neuron a “neural report of time” (NRT, the moment of time the neuron reports as the delay to expected reward, see Methods), we asked if timing activity to a given reward delay similarly emerges in the head-fixed mouse preparation. Should reward timing responses emerge to visual cues predicting a given delay, the central tendency of those cues’ NRT distributions should correspond to that delay. Indeed, we find that the central tendencies for the NRT distributions accord well with the conditioned intervals and are significantly different for short and long delay sessions (Z = −4.95, p = 7.49 × 10^−7^, Wilcoxon rank-sum test – Figure 3D). Furthermore, the NRTs calculated from the cross-validated responses described above also show similar significant changes in distributions (i.e., shorter for the short delay) across the range of values for *k* (all Z’s ≤ −3.43, all p’s ≤ 5.97 ×, 10^−4^ Wilcoxon rank-sum test). This shifting of the NRT distributions across conditioned intervals is not explained by differences in licking behavior as licking alone has no effect on recorded neural activity (p = 0.198 Wilcoxon Sign Rank Test) and an animal’s within-session licking behavior did not influence the calculated NRT (χ^2^(2, 1120) = 1.91, p = 0.385, Kruskal-Wallis Test, Figure 3, Supplemental Figure 1).

In previous reports of reward timing, neurons showed “cue dominance” (i.e., expressing reward timing activity to one, but not both cues) when the two cues are paired with different delays. Here we find that when the two cues predict a reward at the same delay that neurons can express such cue-specificity in reward timing, but that there is an increase in neurons with reward timing to both cues. Specifically, we find that of the 253 neurons that have reward timing, 157 (62%) express reward timing to both cues and do so with notable similarity across cues (Figure 3, Supplemental Figure 2). The remaining 38% express reward timing to one, but not the other cue, despite the cues foretelling of the same magnitude and delay to reward. Additionally, we find that neurons have stable representations of time across days. Using a previously-described statistic based on waveform shape (the J3 statistic, (Moran and Katz, 2014)) we defined 100 pairs of putative repeatedly-recorded neurons across two consecutive sessions. Of the 77 pairs of reward timing responses across days, we find that 45 (58%) have notably similar reward timing responses across days (Figure 3, Supplemental Figure 3).

Together, these data demonstrate that mouse V1 neurons are able to express reward timing activity following associative learning. Having established mouse V1 as a locus for such timing activity, we sought to investigate how inhibitory interneurons express and aid in the expression of this reward timing activity. Specifically, we turned to recent theoretical work which proposes a manner by which neurons in primary visual cortex could create such heterogeneous representations of time (Huertas et al., 2015).

### V1 Neurons Represent Reward Timing in a Manner Consistent with a Theorized Network Architecture

Recent computational work posits that a simple network motif can produce reward timing activity with the three known response forms (Huertas et al., 2015). This network motif is derived from a recurrent network of excitatory cells with broad and sparse inhibition; it contains one population of inhibitory cells and three populations of excitatory cells which differ based on levels of recurrent excitation, non-recurrent excitation, and inhibition (schematized in Figure 1C). Two experimentally tractable implications of this network motif are: (1) inhibitory interneurons should represent reward timing predominantly as the sustained increase form and (2) neurons that are inhibited by interneurons should represent time predominantly as sustained decrease or peak forms. Here we test these predictions using mice which selectively express channelrhodopsin (ChR2) exclusively in one of three major interneuron subtypes: those expressing parvalbumin (PV), those expressing somatostatin (SOM), and those expressing vasoactive intestinal polypeptide (VIP, see Methods).

By investigating the ability of each interneuron subtype to fulfill the model’s implications, we are able to determine how known interneuron diversity intersects with the proposed network architecture. With selective ChR2 expression we are able to optogenetically identify interneurons within our recorded population (Figure 4A-4B, see Methods). We identified 35/185 (18.9%) PV+ neurons, 15/361 (4.2%) SOM+ interneurons, and 0/203 (0%) VIP+ interneurons (example cells shown in Figure 4B). These proportions match the expected relative distribution given our recording depth (Tremblay et al., 2016). Additionally, a control cohort of animals (not expressing ChR2 in any cell population) resulted in 0/247 (0%) recordings returned as expressing ChR2 from these wildtype animals.

**Figure 4:**
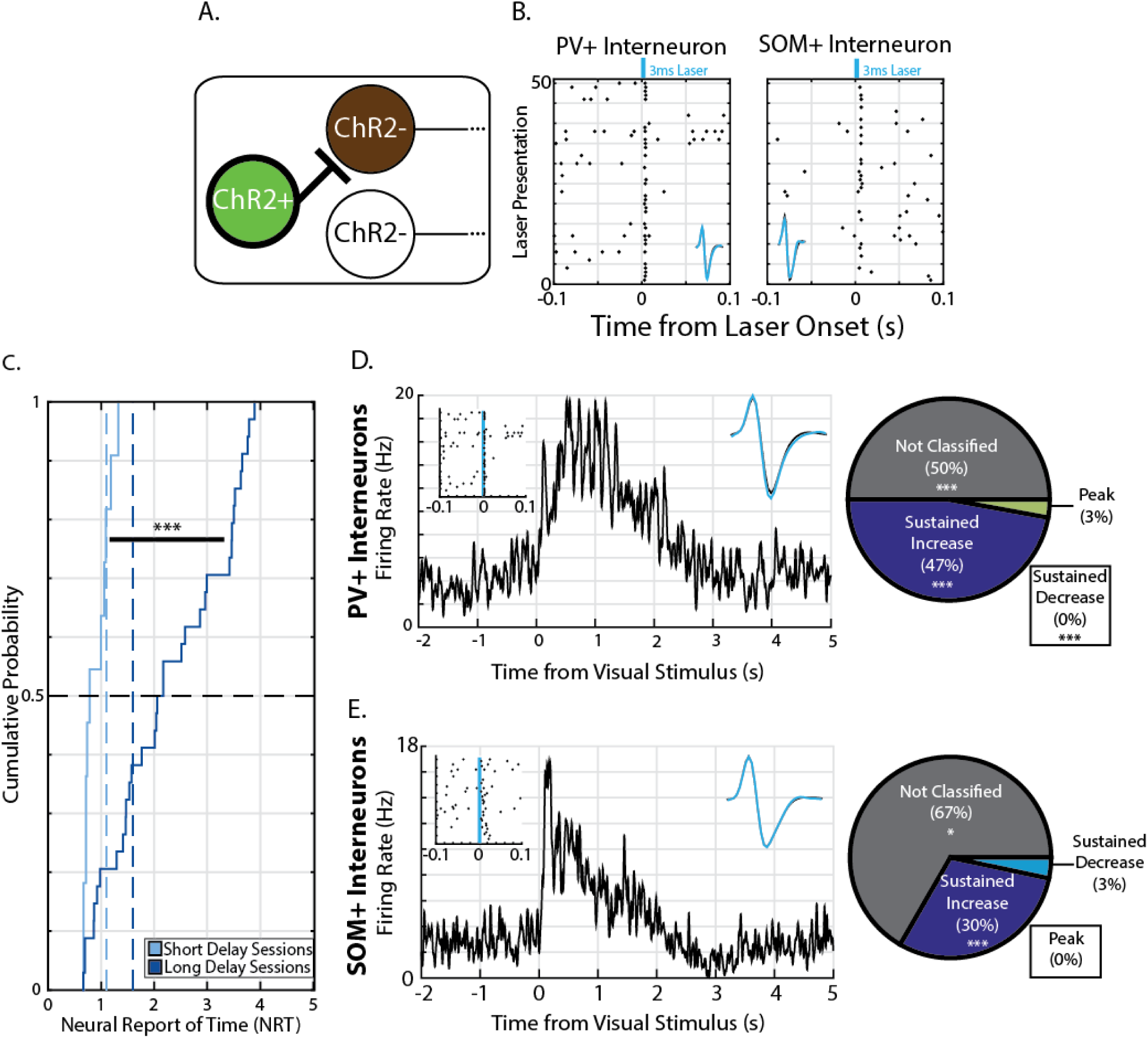
Identified interneurons express reward timing as SI form, consistent with theorized network architecture. (A) Cartoon showing the three types of cells we could possibly record from: ChR2-expressing (ChR2+,green), ChR2-negative which are inhibited by ChR2+ cells (ChR2-, brown), and ChR2-negative that are not inhibited by ChR2+ cells (ChR2-, white). Here we investigated reward timing in neurons expressing ChR2 as indicated with the bold lines around the ChR2+ population. (B) Two example raster plots of spikes recorded during “optotagging”. Spikes are black dots plotted with respect to the time of a 3ms laser stimulus; this activity is recorded from a putative PV+ interneuron (left) and a putative SOM+ interneuron (right) and insets show average waveforms during laser stimulation (cyan) and average waveforms during spontaneous activity (black). (C) Cumulative distribution plots of NRTs calculated from identified interneurons during short delay sessions (light blue) or long delay sessions (dark blue). Vertical, dashed lines represent time of expected reward and horizontal, dashed line indicates median of the distributions. (D) Left: Example PV+ interneuron expresses reward timing as the Sustained Increase form. Inset in top left shows raster plot of spikes during optotagging and inset in top right shows average laser-evoked waveform and average spontaneous waveform (conventions as in B). Right: Pie chart showing proportions of responses from all PV+ interneurons which express reward timing and in what forms. (E) Same as E, but for SOM+ interneurons. * - p < 0.05,*** - p < 0.001, Wilcoxon Rank-Sum test (C) or Bootstrap (D -E).

As we optogenetically identified PV+ and SOM+ interneurons, we were then able to ascertain their reward timing capabilities. We first determined their ability to produce representations of time and found that their NRTs shift across conditioned intervals (Z = −3.605, p = 3.117 × 10^−4^, Wilcoxon rank-sum test; Figure 4C). Again, we verified that these representations of time are unlikely to be explained by licking behavior as licking, by itself, had no significant effect on ongoing spiking activity (p = 0.355, Wilcoxon Sign Rank Test; Figure 4, Supplemental Figure 1). Having demonstrated that identified interneurons are expressing reward timing, we then asked what the distribution of reward timing forms are for the subpopulation of interneurons. We found that 1) PV+ interneurons are significantly more likely to represent the time interval than non-identified counterparts, and, that 2) they are significantly more likely to represent time as a sustained increase of activity (p = 8.10 × 10^−12^, p = 6.33 × 10^−25^, respectively, bootstrap – Figure 4D). We then asked the same of SOM+ interneurons and found that, again, they are more likely to express reward timing and are significantly more likely to represent time as a sustained increase of activity (p = 0.018, p = 3.61 × 10^−4^, respectively, bootstrap – Figure 4E). Additionally, although we did not optogenetically identify excitatory cells, we have identified a subpopulation of putative pyramidal cells using waveform shape. We did so, specifically, by using the spike width of a waveform to define a population of recorded cells as wide-spiking (Barthó et al., 2004). To determine the reward timing expression of putative pyramidal cells, we looked at neurons within the top quartile of the spike width distribution. As expected, we find that these neurons express reward timing in all forms (Figure 4, Supplemental Figure 2). These data are in accordance with the proposed network architecture which suggests that inhibitory interneurons should represent the time interval as the sustained increase form.

An additional component of the theorized network architecture is that neurons whose spiking is inhibited by inhibitory neurons should express reward timing as either the sustained decrease or peak form. To investigate this prediction, we defined cells as “suppressed” by presenting extended laser stimuli (100ms) and recording responses (see Methods). Consistent with this prediction, we found that neurons which are inhibited by PV+ activation are significantly more likely to represent the time interval as the sustained decrease form (p = 1.85 × 10^−4^, bootstrap; Figure 5A). However, neurons that are inhibited by SOM+ activation were less likely to express reward timing (i.e., were more likely to be not classified, p = 1.61 × 10^−5^, bootstrap) and, contrary to the model’s prediction, were significantly less likely to be classified as sustained decrease or peak (p = 7.49 × 10^−5^ and p = 0.02, respectively, bootstrap; Figure 5B). Additionally, we find that neurons that are inhibited by VIP+ activation are significantly more likely to express reward timing (p = 0.018, bootstrap) and have a significant enrichment of the sustained increase form (p = 0.014, bootstrap; Figure 5C). Together, these results show that PV+ interneurons fulfill the expectations of the theorized interneuron population and provide evidence in favor of the proposed network architecture wherein the reward timing forms arise due to the connectivity among excitatory and inhibitory neurons within V1. In addition, these results also point to functional distinctions of interneuron subtypes in the production of reward timing activity.

**Figure 5:**
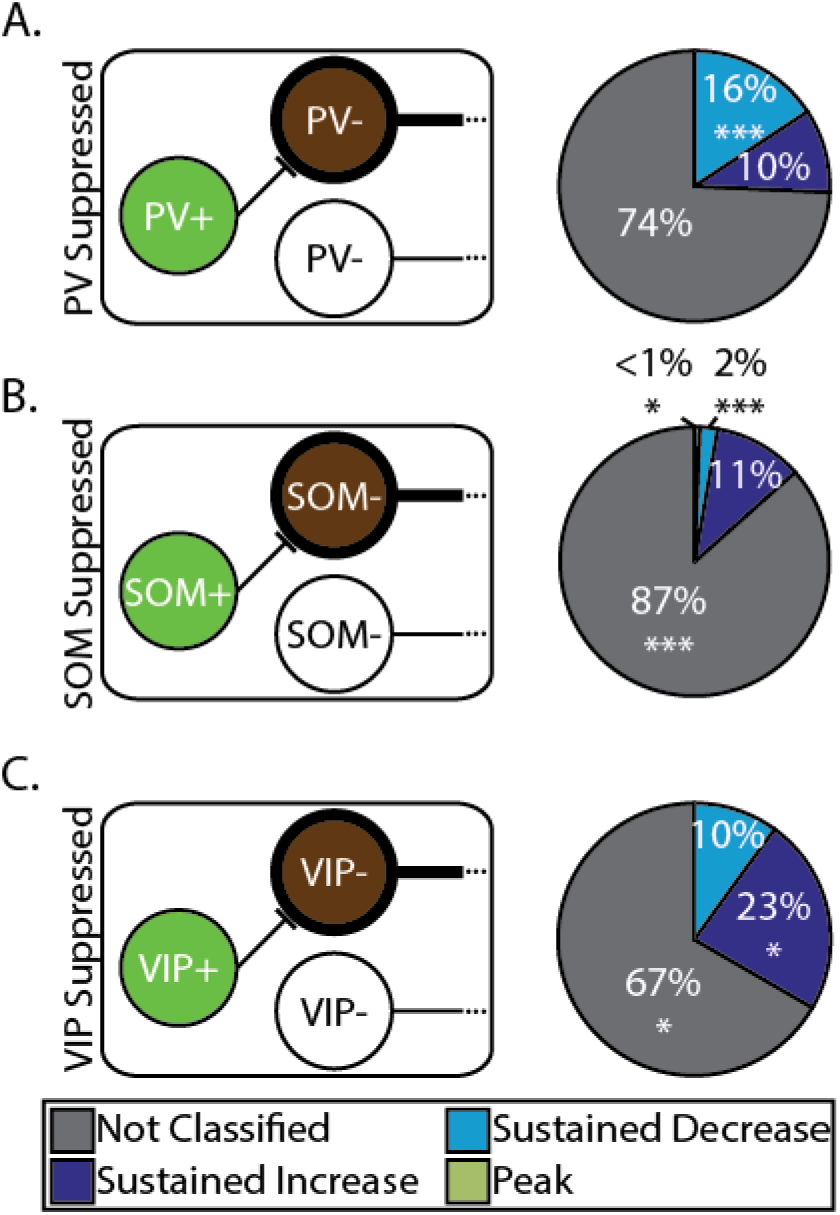
Neurons inhibited by different interneuron subtypes express reward timing forms in varying proportions. (A) Left: Cartoon schematizing specific PV-neurons of interest (i.e., those inhibited by PV+ interneurons); note that though cartoon shows monosynaptic inhibition, we are unable to address the mono- or polysynaptic nature of observed inhibition. Right: Pie chart showing proportion of responses which express reward timing and in what forms. (B) Same as A but for SOM-neurons which are inhibited by SOM+ activation. (C) Same as A but for VIP­ neurons which are inhibited by VIP+ activation.* - p < 0.05, ** - p < 0.01, *** - p < 0.001, bootstrap.

## Discussion

Reward timing in the primary visual cortex is a network phenomenon that requires coordinated activity of various cell types. Here we have shown that reward timing exists in three forms in V1 of head-fixed mice; that reward timing activity is expressed within identified interneuron subpopulations; and, that neurons that are inhibited by different interneuron subpopulations differ in their expression of reward timing. These findings are consistent with a theorized network architecture.

### Reward Timing in the Primary Visual Cortex of Head-Fixed Mice

Head-fixed mice were trained to associate a visual stimulus with a delayed reward (Figure 2) and V1 neurons reflected this learned association in one of three forms (Figure 3). These results replicate previous reward timing reports in the primary visual cortex of freely-moving rats (Shuler and Bear, 2006; Chubykin et al., 2013) and mice (Liu et al., 2015), extend these reports to the head-fixed preparation, and add to reports of non-sensory representations within V1 (Ji and Wilson, 2007; Poort et al., 2015; Fiser et al., 2016; Pakan et al., 2018). As other sensory areas express altered representations following associative learning (McGann, 2015), our understanding of V1 reward timing allows for greater insight into how cortical circuits, generally, can create predictions of future events.

Though we contend that the production of reward timing in V1 is the result of interactions among cells within it, might it be that V1 is reflecting some non-specific global input signal (e.g., arousal or attention)? Our data argue that this alternate explanation is unlikely to be the case. First, a substantial fraction of neurons with reward timing show “cue dominance” (i.e., express reward timing to one, but not both cues, Figure 3 Supplemental Figure 2). Such specificity in reward timing is difficult to explain if V1 neurons were reflecting some non-specific, global signal. Second, we find that the expression of reward timing is unaffected by how engaged an animal is in our task, as indicated by its licking behavior (a measure known to co-vary with other measures of arousal (Lee and Margolis, 2016)). The dissociation between licking and reward timing, then, is not consistent with a global signal being the cause of V1 reward timing activity.

### V1 Neurons Express Reward Timing in a Manner Consistent with the Core Network Architecture

Recent computational work has theorized a manner by which a network of cells can produce reward timing activity in the various forms observed (Huertas et al., 2015). The results of this formal model suggest that the heterogeneity of reward timing forms can be captured by a “core network architecture” where the relative amount of inhibition, recurrent excitation, and non-recurrent excitation differs according to a simple motif (Figure 1C). Here we sought to address two key implications of this model to determine potential biological validity of the proposed network architecture: 1) inhibitory interneurons should reflect reward timing predominantly as the sustained increase form and 2) neurons inhibited by interneurons should express reward timing predominantly as the sustained decrease or peak forms. Using selective expression of channelrhodopsin-2 (ChR2) in interneuron subpopulations, we are able to define these two populations (interneuron and suppressed) outside of behavioral conditioning and probe the reward timing expression in such populations. A simplifying assumption of the model is that all interneurons behave in a similar manner; however, it is known that there are various different interneuron subpopulations within V1 and that they have been shown to have different functional roles when the network represents sensory information (Atallah et al., 2012; Lee et al., 2012; Wilson et al., 2012; Kvitsiani et al., 2013). By selectively expressing ChR2 in specific interneuron subpopulations, we are able to address the model’s implications and determine how these activity patterns intersect with the various interneuron subtypes.

Here we identify putative PV+ and SOM+ interneurons (Figure 7B-D) and find that PV+ interneurons adhere to the model implications. They produce reward timing predominantly as the sustained increase form (Figure 4D), and neurons that they inhibit produce reward timing with an enrichment of the sustained decrease form (Figure 5A). These results can be contrasted with SOM+ interneurons which, while expressing reward timing predominantly with the sustained increase form (Figure 4E), largely do not inhibit neurons which express reward timing (Figure 5B). Finally, the manner by which VIP+ interneurons express reward timing is unknown, but we have shown that those neurons inhibited by VIP+ activation express reward timing predominantly with the sustained increase form (Figure 5C). These results can be understood when known connectivity is incorporated into the network architecture, as discussed below.

Inhibitory interneurons are known to have distinct connectivity among other interneurons and pyramidal cells (Pfeffer et al., 2013; Tremblay et al., 2016). For example, it is known that VIP+ interneurons predominantly innervate other inhibitory interneurons (Pfeffer et al., 2013; Tremblay et al., 2016). As such, it follows that the neurons we defined as inhibited by VIP+ interneurons express reward timing in a manner similar to PV+ and SOM+ interneurons (i.e., with an enrichment of the sustained increase form). Additionally, the finding that PV+ and SOM+ interneurons produce reward timing in a similar manner may be surprising as they are thought to perform different functions in stimulus representation (Atallah et al., 2012; Lee et al., 2012; Wilson et al., 2012). However, according to the theorized network architecture, interneurons express reward timing as the sustained increase form because they receive input from the excitatory population of sustained increase neurons (Figure 1C). Cortical interneurons are known to receive convergent input from local excitatory cells (Bock et al., 2011; Fino and Yuste, 2011; Hofer et al., 2011; Packer and Yuste, 2011). Specifically, a population of deep layer pyramidal cells has been shown to target both PV+ and SOM+ interneurons (West et al., 2006). Perhaps the similarity in reward timing expression in PV+ and SOM+ populations arises from similar pyramidal cell input to these interneurons. Additionally, if this input is shared with VIP+ interneurons, it would posit that VIP+ interneurons could also express reward timing as the sustained increase form. Thus, functional differences would be borne out in downstream neurons (e.g., those neurons whose activity is suppressed by interneurons, Figure 5). Finally, when comparing between PV+ and SOM+ interneurons, PV+ interneurons act in a manner more consistent with the inhibitory population proposed in the core network architecture. This also can be understood when one considers that PV+ interneurons are the most abundant type of interneuron and provide the majority of inhibition to pyramidal cells (Markram et al., 2004; Tremblay et al., 2016). Taken together, we are able to overlay known connectivity patterns with the data shown here to hypothesize amendments to the core network architecture (Figure 6).

**Figure 6:**
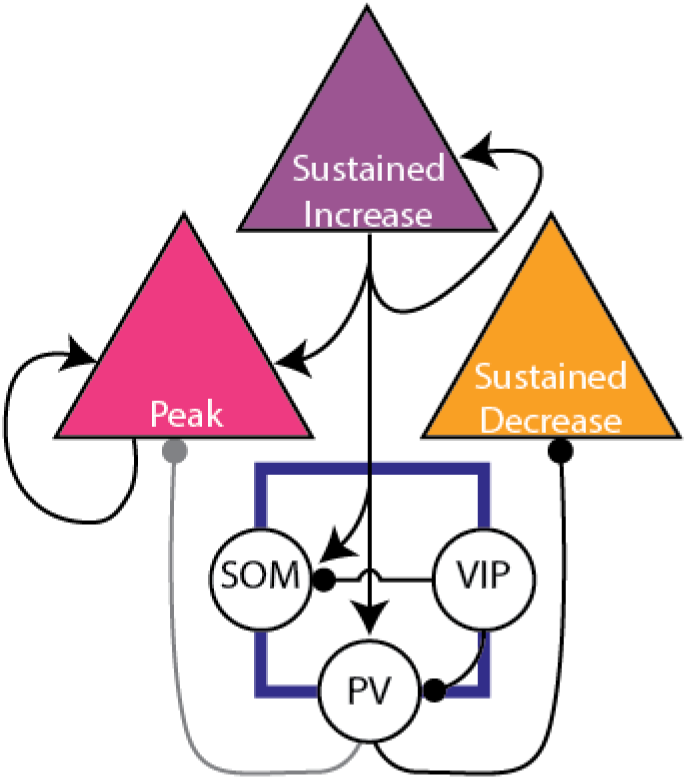
Predictions for core network architecture. The network architecture adapted from Figure 1 showing a potential manner by which simulated inhibitory neurons can be divided based on reward timing responses and reported connectivity. The gray line connecting PV+ interneurons is purported to exist given computational work; while not evidenced in the data presented here, it cannot be ruled out. In the current proposed architecture, PV+ interneurons fulfill expectations of the theorized inhibitory population as they produce reward timing with the SI form and inhibit neurons expressing reward timing with the SD form.

### Concluding Remarks

Reward timing in the primary visual cortex, like many complex responses in the brain, requires the coordinated activity of many cells. Here we have enriched the phenomenological characterization of reward timing, extended that work to incorporate the heterogeneity of cell types within V1, and provided a greater comprehension of how V1 produces reward timing activity through its circuitry. We now better understand how the various cell types come together to produce such a representation of time between a cue and a reward.

## Materials and Methods

### Experimental Design and Statistical Analyses

All procedures performed on animals were in accordance with the US NIH Guide for the Care and Use of Laboratory Animals and were approved by the Animal Care and Use Committee at the Johns Hopkins University School of Medicine. Male mice (N = 14, between 2 and 6 months old) were used in this study. For this study, we used four genetic backgrounds which differ in their expression of the light-activated cation channel, channelrhodopsin-2 (ChR2). One cohort expressed no ChR2 (wildtype or WT, n = 3 animals), one cohort expressed ChR2 in parvalbumin-positive interneurons (PV-ChR2, n = 4 animals), one cohort expressed ChR2 in somatostatin-positive interneurons (SOM-ChR2, n = 4 animals), and one cohort expressed ChR2 in vasoactive-intestinal-polypeptide-positive interneurons (VIP-ChR2, n = 3 animals). The WT cohort was composed of C57BL/6 mice (Strain Code: 027, Charles River Laboratories). ChR2-expressing mice were derived by selectively breeding the following genetic lines: a parvalbumin-Cre recombinase line (PV-Cre; 008069, Jackson Laboratory, (Hippenmeyer et al., 2005)), a somatostatin-Cre recombinase line (SOM-Cre; 013044, Jackson Laboratory, (Taniguchi et al., 2011)), a vasoactive-intestinal-polypeptide-Cre recombinase line (VIP-Cre; 010908, Jackson Laboratory, (Taniguchi et al., 2011)), and a loxP-STOP-loxP-channelrhodopsin-2-eYFP Cre-dependent line (ChR2-eYFP, Ai32; 012569, Jackson Laboratory, (Madisen et al., 2010)). The mice of these crosses were on mixed backgrounds composed primarily of C57BL/6 and CD-1. All mice from all cohorts underwent identical training and training occurred in the light cycle during a 12h light/dark schedule (lights provided between 0700 and 1900).

Non-parametric tests (e.g., Kruskal-Wallis test and Wilcoxon Rank-Sum test) are used throughout the text with an alpha of 0.05 unless otherwise noted. Other statistical analyses include the use of the Stimulus Associated Latency Test (SALT) as described previously (Kvitsiani et al., 2013) and bootstrap analyses; both methods are described in more detail below.

### Surgical Procedures

Surgical procedures were performed under aseptic conditions and were in accordance with the Animal Care and Use Committee at the Johns Hopkins University School of Medicine. Animals underwent two surgeries spaced at least two weeks apart from one another. Prior to either surgery, mice were anesthetized using a cocktail of ketamine (Ketaset, 80mg/kg) and xylazine (Anased, 10 mg/kg) and eyes were covered with ophthalmic ointment (Puralube). The first surgery was performed to affix a head-restraint bar to the animal’s skull for training purposes and to mark sites for future craniotomies. In the first surgery, the hair covering the skull was removed (Nair), the skin cleaned with alternating 70% ethanol and iodine, then the skin was cut away. Following this, the periosteum was removed and the skull cleaned with alternating 70% ethanol and hydrogen peroxide, then the skull was dried with canned air. A total of four sites were marked for future craniotomies: two for ground screws (arbitrarily marked over the anterior parietal bone) and two for primary visual cortex (measured as 3mm lateral to lambda, bilaterally). Sites for future craniotomies were covered in a silicone elastiomer (Smooth-On Body Double) and a head-post was affixed to the anterior portion of the mouse’s skull with super glue (Loctite 454). The remaining bone was covered in super glue. A second surgery was performed to implant recording electrodes. Briefly, small craniotomies were performed using a dental drill for ground screws and screws were implanted into sites. Next, craniotomies were performed over V1, the dura cleaned with sterile paper points, and electrodes were brought to the surface of the brain, then implanted 500μm below the cortical surface in accordance with stereotaxic measurements of V1 (Franklin and Paxinos, 2008). Wires were covered in sterile ophthalmic ointment (Puralube) and encased in dental cement (Orthojet). Ground screws and ground wires were connected and a headcap was built of dental cement.

### Behavioral Task Design

Prior to electrode implantation (between the first and second surgeries), animals were habituated to head-fixation over the course of 2-3 days, and then were trained that a visual stimulus predicted a water reward at a fixed delay for 2-3 weeks. Visual stimuli were full-field retinal flashes delivered monocularly to the left (Cue 1) or right eye (Cue 2) via head-mounted goggles. These goggles are custom made and consist of a miniature LED glued to the back of a translucent, plastic hemidome. Licks were recorded on a lickometer via an infrared beam break (IslandMotion); experiments were controlled through an Arduino Mega microcontroller board (Arduino) and events were recorded with Neuralynx. In every session, trials were separated by an inter-trial interval (ITI, between 3 and 8 seconds, uniformly distributed). In order to initiate the next trial (and exit the ITI), animals had to cease licking for a random interval during the later portion of the ITI (deemed a “lick lockout”). This lick lockout period was the same across conditioned delays and was used to discourage non-stimulus-evoked licking, as licks within this period caused the timer to restart and, thus, a longer ITI. Upon trial initiation, a monocular visual stimulus was either delivered (CS trials) or withheld (Sham trials), followed by a delay window. CS’s were visual stimuli which lasted 100ms and were delivered, with equal probability, to the left or right eye. The delay to reward was the same for both CS’s within a session and was held constant for several consecutive sessions as either the short (1s) or the long (1.5s) delay. Sessions conditioned with the short delay constitute the “short delay sessions”; those with the long delay, the “long delay sessions”. CS trials were further divided into “paired” and “catch” trials; paired trials being trials in which a small water reward (∼2μL) became available following the delay period, provided that the animal made at least one lick on the lick port within the delay. Catch trials, however, were trials in which the reward was withheld regardless of behavior. Licks were never rewarded during Sham trials. At the conclusion of the delay window on both CS and Sham trials, the animal re-entered the ITI. Trials in which the animal licked during the delay window are defined as “Hit” trials and trials in which the animal did not lick during the delay window are defined as “Miss” trials. Unless otherwise noted, data presented here are from Catch+Hit trials (i.e., trials in which the animal received a visual stimulus, licked during the delay window, and did not receive a water reward at the end of that delay).

The relative proportion of paired/catch trials and sham trials was systematically varied across behavioral shaping as well as the requirement to lick within the delay window. In the final form of the task (and in all sessions reported here), 80% of trials were CS trials (with equal probability of being paired or catch), with the remaining trials being Sham trials.

### Behavioral Measurements

The timing of individual licks was recorded using a lickometer (IslandMotion) and were recorded simultaneously with neural data. During the task, animals tended to make one lick bout following delivery of the CS; the timing of this bout is quantified as the time of the first lick within the bout, the time of the last lick within the bout, and the mean time between these two licks (“Bout Midpoint”).

### Electrophysiology

Neural activity was recorded bilaterally from primary visual cortex using custom-built recording electrodes. Per recording electrode, 16 channels of neural data were recorded at a sampling rate of 32,556 Hz through commercial hardware (Neuralynx). Neurons were offline identified through manual, 3D cluster-cutting methods through commercial software (Offline Sorter, Plexon). Electrodes were composed of a connector with 16 recording channels and two ground wires (Omnetics). Bundles were cut at a ∼45° bias to allow for sampling across a depth of approximately 250µm. In order to optogenetically identify interneuron subtypes, an optic fiber—composed of a 200µm core diameter glass multimode fiber (ThorLabs) and a 1.25mm ceramic stick ferrule (Precision Fiber Products)—was glued next to the wire bundle such that the tip of the optic fiber was abutted next to the majority of the electrode tips of the bundle (<200µm tip-to-tip distance with some wires above optic fiber tip, some next to, and some below optic fiber tip). A schematic showing the recording strategy can be found in Figure 1A.

### Neural Data Analysis

The following neural data analyses were performed using custom scripts and functions in MATLAB (Mathworks).

#### Reward Timing Classification

The form with which a neuron expressed reward timing was determined using manual classification in a blinded fashion. Specifically, a neuron was randomly selected from a random session. Then, a peri-stimulus time histogram (PSTH) calculated from trials that were either Cue 1 Catch+Hit trials, Cue 2 Catch+Hit trials, or Sham+Hit trials was randomly presented to an experimenter (KJM). This PSTH was then classified as “Not Classified” (NC), “Sustained Increase” (SI), “Sustained Decrease” (SD), or “Peak” (PK). The remaining PSTH’s were presented, followed by the remaining neurons. These classifications were performed without knowledge of animal identity, recording session, or delay time.

#### k-Nearest Neighbors Classification

We sought to cross-validate the human classification of reward timing neurons. To do so, we implemented a *k*-Nearest Neighbors (kNN) classifier. Briefly, kNN takes classified data as a “training example” to then determine the identity of unclassified “query points” based on the proximity of query points to classified training example points. Identity of the query point is defined as a plurality vote of its *k* nearest neighbors in the training example. In our case, we first split data from reward timing neurons into two halves: neural activity from even trials and neural activity from odd trials. Then, we normalized neural activity using the area under the ROC curve (AUC, see below) and used principal components analysis (PCA) for dimensionality reduction. Specifically, we reduced the normalized firing activity from even trials to the first eight principal components which explained >85% of the variance within the neural activity; the projection in eight dimensions and human-classified identity of the responses recorded in even trials served as the training example for the kNN classifier. Then, data from the odd trials were projected into the 8-dimension subspace (acting as the query points) and were classified across a range of *k*. Specifically, we varied the number of neighbors between 1 and 65; to avoid ties, we only used an odd number of neighbors in our classification.

#### Neural Report of Time Classification

To attribute a time in which neurons with reward timing activity reported the expected delay to reward, we calculated the Neural Report of Time (NRT). The NRT is the moment taken as the time which neurons return to a baseline level of activity, for SI and SD response forms, or the time of maximum firing rate from baseline (after the visual-evoked response), for the PK response form.

To calculate such a time, neural activity was normalized to the baseline firing rate by calculating the area under the ROC curve (AUC) using a sliding 100ms window (Cohen et al., 2015; Sadacca et al., 2018). An AUC value of 0.5 means that the ideal observer would be at chance level to tell apart two distributions and values above or below 0.5 reflect greater dissimilarity among two distributions. For our purposes, we found the AUC value between the distribution of spike counts from a 100ms window of baseline pre-stimulus activity, and a given 100ms of spiking activity across all trials of the same type (e.g., Paired, Cue 1 trials; Catch, Cue 1 trials; etc.). In this way, we do not rely on the averaging of spike counts in the same way that a PSTH does and thus the resultant value is more robust against a small subset of trials with many spikes or other forms of inter-trial spiking variability. Furthermore, this method normalizes the firing rate to a value bounded by 0 and 1 for every set of trials. As the AUC-normalized firing rate is the magnitude of difference and not the sign of the difference between an AUC value and 0.5 (which determines how dissimilar two distributions are), we found the absolute value of the difference between the AUC vector and a value of 0.5. In doing so, neurons with sustained activation or suppression (SI or SD neurons, respectively) could be treated with the same algorithm to calculate an NRT. We operationally defined a difference threshold of 0.15 (true AUC value of 0.35 or 0.65), and, using this threshold, we then defined the NRT as the first moment in time when the AUC difference vector fell below the threshold for at least 100ms. For classified PK neurons, the NRT was defined as the time of the maximum of this AUC difference vector. To avoid conflating reward timing responses with general visual responses, we set a minimum value for valid NRTs as 0.5s after stimulus offset. Though only Catch+Hit and Sham+Hit trials were classified, we were able to use the Catch classifications to calculate a response’s NRT in two other trial types: Paired+Hit and CS+Miss. The algorithm for calculating these NRTs was identical across trial types.

#### Calculation of ΔSpikes

This value is used to determine the average change in spike rate based on an animal’s first lick in Sham+Hit trials. It is defined as follows:

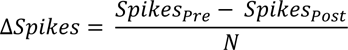

Where *Spikes_Pre_* is the number of spikes in the 100ms preceding the first lick within a Sham+Hit trial, *Spikes_Post_* is the number of spikes in the 100ms following the first lick within a Sham+Hit trial, and *N* is the number of trials of Sham+Hit trials within the session.

#### Calculation of the J3 Statistic

This statistic was developed to determine whether neurons are the same from one recording session to the next (Moran and Katz, 2014). First the waveforms of all spikes recorded from two recordings are projected onto reduced dimensions using PCA. Then, values are calculated as follows:

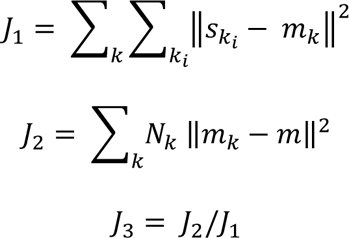

Where 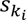 is the projection in two dimensions of spike *i* in session *k*, *m_k_* is the mean vector of all spikes (*N_k_*) from the *k*^th^ session, *m* is the overall point mean of the projection, and ||·|| represents the Euclidean Distance. In essence, the J3 value is a ratio between the Euclidean distance between each spike’s waveform and the center of the cluster of all other spikes’ waveforms from that neuron to the distance between the two clusters (i.e., a ratio of the inter- and intra-cluster distance). J3 is maximal when two recordings are tightly packed and far away from one another in PC space; this reflects that two recordings are unique from one another. However, we utilized this statistic to determine whether a neuron recorded on one day was the same as a recording made on the same channel the subsequent day. To do so, we defined a J3 threshold by finding all “within” J3 values (that is, the J3 value between the first third of the recording’s spikes and the last third of the recording’s spikes). The threshold was defined as the 95^th^ percentile of this distribution. That is to say, any neurons which were recorded from the same animal and on the same channel which had a J3 value that was less than this threshold was deemed the same.

#### Similarity Measurements of Reward Timing Responses

We sought to assess the similarity of reward timing responses of a given neuron across the two CS’s to assess the consistency of reward timing responses when different cues predicted the same reward occurring at the same delay. Furthermore, where possible, we sought to assess the stability of a neuron’s reward timing response to the same stimulus across sessions. Reward timing responses of a neuron could differ (or persist) between cues or across sessions in their presence, form, timing, and shape. For instance, within a session, reward timing responses may be present within a given neuron to both cues, exhibit the same response form (e.g., SI) with an overall similar response shape, and report nominally similar NRT’s. Additionally, neurons can express similar responses to the same stimulus across days. To determine how similar these responses are, we first calculated the concordance of reward timing forms (for example, how often a SI cell expresses reward timing as SI for the opposite CS or on a following day). Among the responses which are concordant, we then determined the similarity in the neuron’s report of time by calculating the absolute difference in NRT’s. Finally, within these responses, we quantified the similarity in shape by calculating the Euclidean distance between the evoked responses. These values were compared with a shuffled control distribution. Shuffling distributions were calculated by shuffling across neurons that expressed reward timing in the same form for the same conditioned interval.

### Neuron Identification

Mice in this study (with the exception of the WT cohort) expressed channelrhodopsin-2 in one of three interneuron populations: PV+, SOM+, and VIP+ interneurons (Figure 1B). This selective expression allows us to determine how the diversity of V1 inhibitory interneurons intersects with the theorized network architecture (as schematized in Figure 1C).

#### Optogenetic Interneuron Identification

Outside of conditioning, brief (1 or 3ms) laser stimuli were randomly delivered to V1 with an inter-pulse interval randomly drawn from a distribution (between 5 and 10 seconds, uniformly distributed) while recording from neurons. To identify putative neurons expressing ChR2, we used the latency to the first spike and the probability that a laser evoked a spike. To determine significant latencies to the first spike, we used the calculated *p*-value from the Stimulus Associated Latency Test (SALT). This test has been previously described (Kvitsiani et al., 2013); briefly, this test compares the latencies to a first spike after a laser stimulus to the latencies to a first spike after arbitrary moments in time without a laser presentation. Specifically, a raster of spiking activity is divided into *N* 10ms bins and the time to a first spike within each bin is recorded. Of the *N* bins created, one bin is the “test bin” and begins with the laser stimulus onset and one other bin is the “baseline bin” (a bin from the pre-laser time period). For all *N* bins, a histogram of first-spike latency is created and a modified Jensen-Shannon divergence is calculated between pairs of these distributions. The divergence between the “baseline bin” and all other non-test bins creates a null distribution against which the divergence between the “baseline” and “test” bin is compared. The resultant *p*-value represents the probability that the divergence between the baseline and test bins falls within the null distribution; we have set a conservative alpha of 0.01 as was used in the first description of the method (Kvitsiani et al., 2013). In this way, neurons which have fast and consistent spikes (i.e., fire quickly and with low jitter) after a laser stimulus will be deemed significant. A caveat to this statistical measure occurs when a neuron has a relatively low baseline firing rate. In such a neuron, due to very low firing rates, random, spontaneous activity occurring within the test window may result in a highly-significant *p*-value. For this reason, we also required a neuron to have an action potential in the window immediately following the laser at least 20% of all laser stimulus presentations.

#### Identification of Pyramidal Cells via Spike Width

In addition to interneuron identification we sought to define a population of putative pyramidal cells. We did so by calculating a neuron’s spike width where the spike width is defined as time difference when the average waveform first crosses 20% of its peak amplitude and last crosses 20% of its valley amplitude. We then set a threshold at the 75^th^ percentile of non-identified interneurons to define a population of putative pyramidal cells.

#### Optogenetic Identification of Suppressed Neurons

Additionally, we were interested in classifying neurons whose responses were inhibited by activating ChR2-expressing interneurons (“suppressed neurons”). Specifically, we sought to classify those neurons that putatively do not express ChR2 (i.e., did not pass one or both of the thresholds set to define interneurons, see above). To determine this, we also presented 100ms laser pulses after the brief laser presentations (with the same inter-pulse interval parameters). We then compared the distribution of spike counts in the 100ms immediately prior to and during laser stimulation with the Wilcoxon signed-rank test (WSRT). If a significant difference was found, we then compared the total number of spikes between these two windows across all presentations. Significantly inhibited neurons are those neurons which passed the WSRT and had fewer spikes during laser presentation than before laser presentation. Although we cannot resolve the exact nature of this inhibition (either mono- or polysynaptic), we are able to assess whether populations affected by inhibitory subtype activation follow predictions of the computational model and whether they reveal functional specialization of various interneurons. Additionally, we have limited our analysis to only those neurons which are inhibited by interneuron activation as neurons which are activated during this stimulation could be activated for one of at least two reasons: (1) they become disinhibited upon activation of interneurons or (2) they express ChR2 but do not pass our statistical thresholds to be defined as expressing ChR2.

#### Bootstrap Procedures

To determine significant changes in the proportion of neurons expressing reward timing in the various forms, we used bootstrap analyses. Specifically, for a given population of “test” neurons (e.g., interneurons or suppressed neurons), we randomly selected a sample of neurons of the same size (with replacement) from all other neurons recorded from animals of the same genotype. We then determined the expression of reward timing in this subsampled distribution and created a bootstrap distribution by repeating the process 1,000 times. *P*-values are the probability that values found in the “test” sample would fall in the bootstrap distribution.

##### Histology

Animals were deeply anesthetized using sodium pentobarbital (200mg/kg, Vedco). After which, animals were transcardially perfused with ice cold phosphate-buffered saline (PBS) followed by ice cold 4% paraformaldehyde (PFA). Brains were immersion fixed overnight in 4% PFA and were transferred to 30% sucrose until sectioning. Brains were sectioned on a cryostat into 60μm slices. Electrode location was verified using Nissl staining, as follows. Sections containing V1 were selected and mounted on gelatin subbed slides and air dried. These slides were then immersed in a solution containing 0.1% Cresyl violet and 1% glacial acetic acid dissolved in water for 5 minutes, followed by a 2-minute wash in distilled water, then by 2 minutes in 50% ethanol, then 2 minutes in 70% ethanol. Stained and washed sections were air dried, immersed in xylenes then coverslipped with Permount Mounting Medium (Electron Microscopy Sciences).

Expression of ChR2 in interneuron subpopulations was verified with immunohistochemistry, as follows. Brain sections containing V1 were selected for immunohistochemistry. On day 1 the sections were washed three times for ten minutes each (3x10 minutes) with PBS then were blocked in 10% normal goat serum (NGS) in PBS + Triton 0.1% to permeabilize and reduce background binding to antibodies for 1h. Sections were then incubated with two primary antibodies overnight at 4°C. Sections for all animals were incubated with a primary GFP antibody to recognize the eYFP tag of the ChR2 (Chicken polyclonal, 1:2000, Aves Labs (Catalog Number: GFP-1020)) and one primary antibody to recognize one of three interneuron markers: PV (rabbit polyclonal, 1:2000, Swant (Catalog Number: PV27)), SOM (rat monoclonal, 1:800, EMD Millipore (Catalog Number: MAB354)), or VIP (rabbit polyclonal, 1:2000, Immunostar (Catalog Number: 20077)). Sections were then washed 3×10 minutes with PBS, then incubated overnight at 4°C with secondary antibodies: Alexa 488 Goat Anti-Chicken (1:500, Jackson ImmunoResearch (Catalog Number: 103-545-155)) and Alexa 568 Goat anti Rabbit (PV and VIP) or Rat (SOM) (1:500, Jackson ImmunoResearch (Catalog Numbers: 111-065-144 and 112-585-143)). Sections were washed with PBS, mounted on glass slides, and coverslipped with Fluoromount-G mounting medium (Electron Microscopy Sciences). To control for unspecific staining, sections were stained in an identical manner except primary antibodies were omitted. Co-expression was as expected given animal’s genotype (Figure 1B).

## Acknowledgements

We wish to thank Harel Shouval, Patricia Janak, Solange Brown, Jeremiah Cohen, Tanya Marton, and Bilal Bari for insightful discussions of this work and comments on the manuscript.

## Grants

Funding for this work was provided by the National Institutes of Health (Grants R01 MH093665 and R01 MH112789 to MGHS) and two training grants (T32 EY007143 and T32 EY017203 to KJM). Microscopy was performed through the NINDS Multiphoton Imaging Core (P30 NS050274).

## Disclosures

The authors declare no competing financial interests.

## Figure Supplements

**Figure 3, Supplemental Figure 1:**
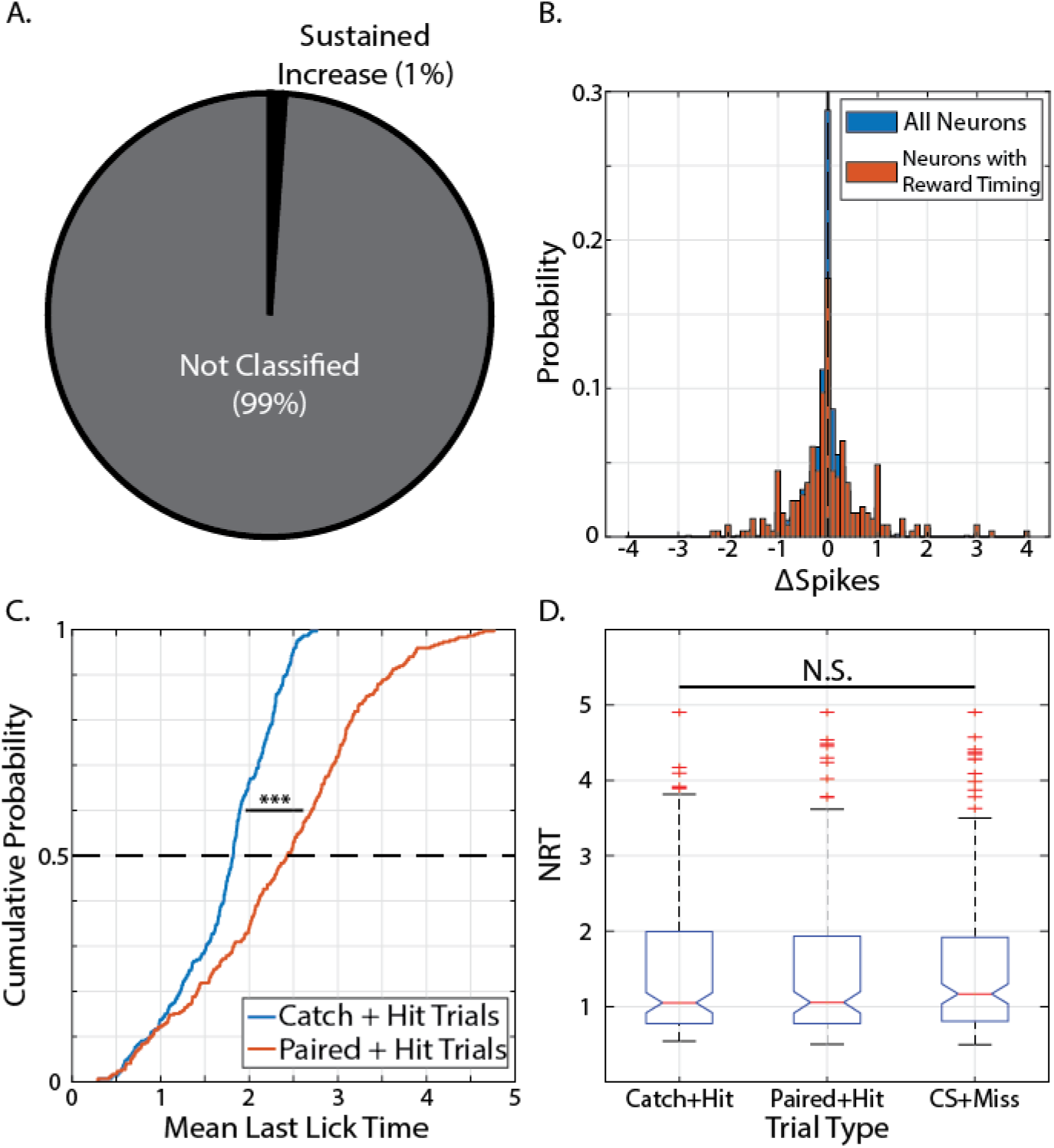
Reward timing activity is not explained by licking behavior. (A) Pie chart showing classification of responses recorded during Sham+Hit trials (trials in which no CS was given, but an animal licked). (B) Distribution of ti.Spikes (a variable which reflects the average change in the number of spikes before and after a first lick, see Methods) for all recorded neurons (blue) and neurons with reward timing responses (orange). Vertical, dashed line represents medians of distributions; neither distribution is significantly different from zero (p’s > 0.05, Wilcoxon Signed-Rank test). (C) Cumulative distribution plots of Mean Last Licks within Catch+Hit trials (blue) and Paired+Hit Trials (orange) show that animals have longer lick bouts for Paired+Hit Trials compared to Catch+Hit trials (Z −13.34, p 1.30 × 10^−40^, Wilcoxon Sign-Rank test). (D) Boxplot of distributions of NRTs calculated from the different trial types from neurons with reward timing responses. Box limits in box plots represent 25th and 75th percentiles, lines correspond to roughly +/-2.7σ. N.S.: Not Significant;***: p < 0.001, Wilcoxon Sign-Rank test.

**Figure 3, Supplemental Figure 2:**
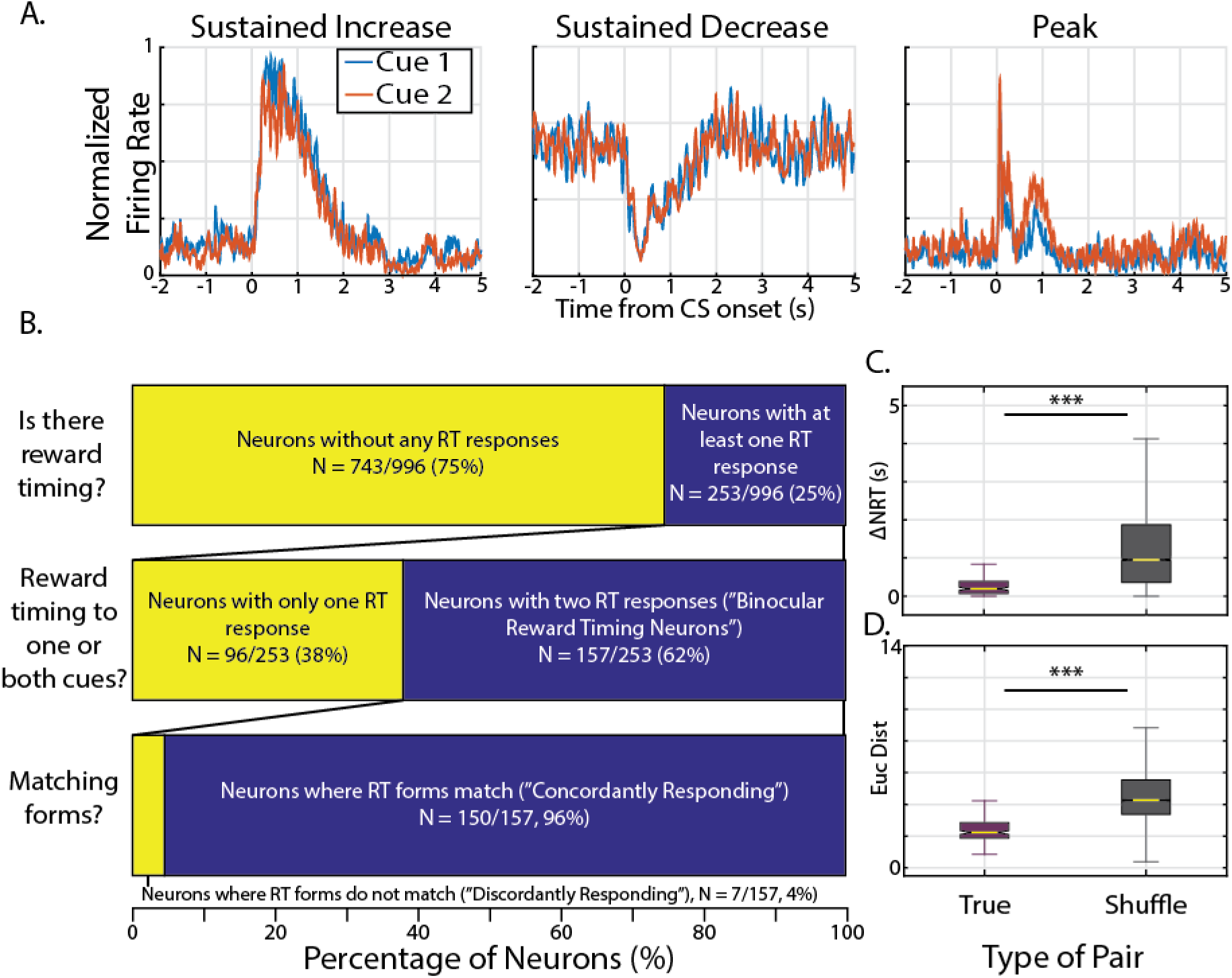
Different cues elicit responses similar in form, timing, and shape within binocular reward timing neurons. (A) Normalized activity from three example neurons which have similar responses to Cue 1 as they do for Cue 2. (B) Bar chart representing the proportion of neurons with reward timing responses (top); of those cells, the proportion of cells with two classified responses (”Binocular Reward Timing Neurons”, middle); and, of the Binocular Reward Timing Neurons, the proportion of neurons which express reward timing as the same form to either CS (”Concordantly Responding”). (C - D) Box plots showing differences in the absolute difference of calculated NRT’s (”LlNRT”, C) and Euclidean distances (D) for “true pairs” (Cue 1 vs Cue 2 of the same neuron) and “shuffle pairs” (Cue 1 vs Cue 2 of neurons with same reward timing form and same conditioned interval). Box limits in box plots represent 25th and 75th percentiles, lines correspond to roughly +/-2.7σ. For demonstration purposes, outliers have been removed from plot.*** in panel C: Z = −12.67, p = 8.60 × 10^−37^, Wilcoxon rank-sum test.*** in panel D: Z −16.97, p 1.39 × 10^−64^.

**Figure 3, Supplemental Figure 3:**
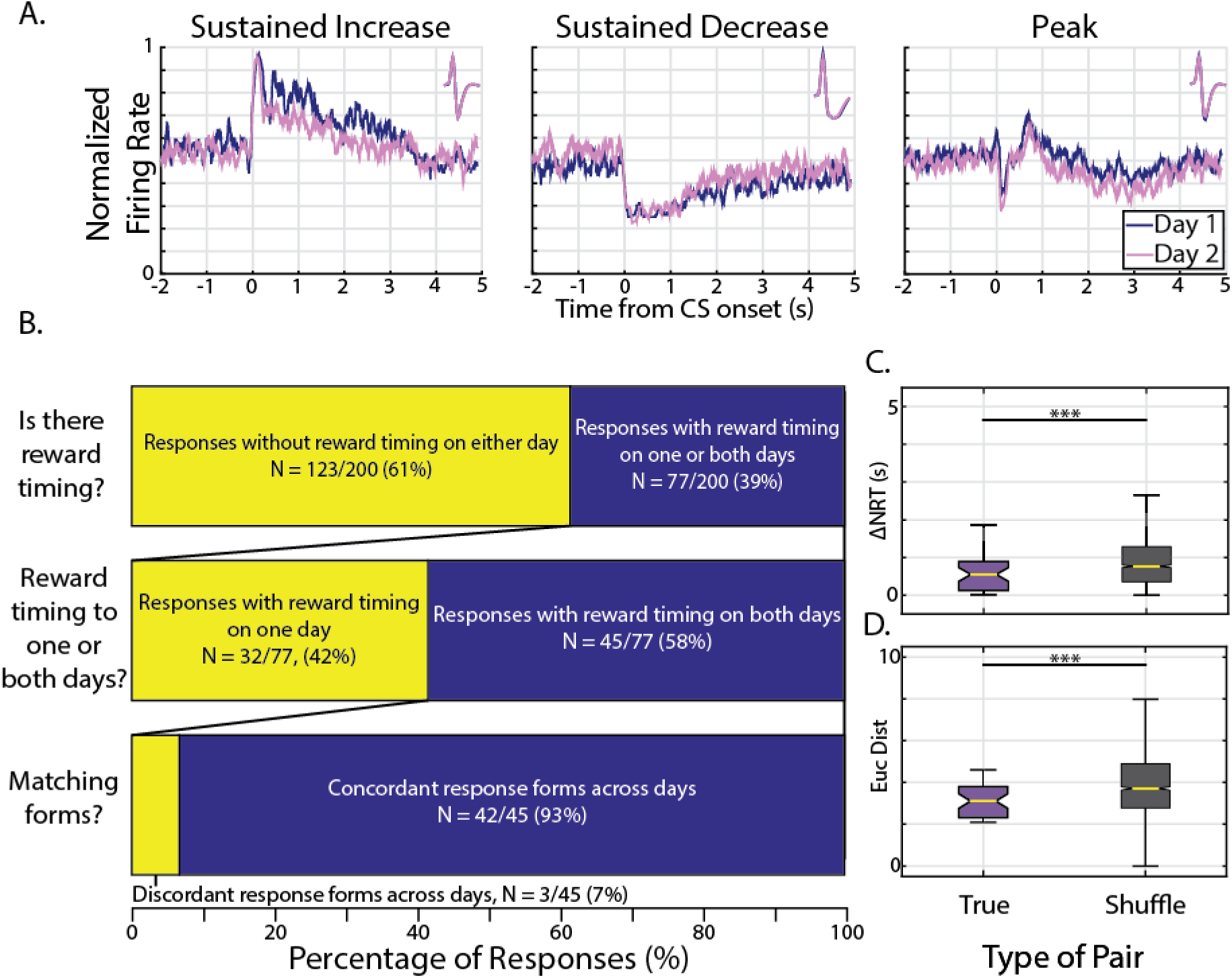
Reward timing is stable in form, timing, and shape across recording sessions. (A) Normalized activity from three example neurons deemed to be the same neuron from different recording sessions which express reward timing similarly on day 1 (purple) and day 2 (pink); average waveforms from entire session shown in insets. (B) Bar chart representing, of all CS responses from repeat neurons, the proportion of responses with reward timing on at least one day (top); of those responses with reward timing on at least one day, the proportion of responses which had reward timing on both days; and of those responses, the proportion of responses which were classified as having the same form (”Concordant Responses”). (C - D) Box plots showing differences in the absolute difference of calculated NRT’s (”ΔNRT”, C) and Euclidean distances (D) for “true pairs” (Day 1 vs Day 2 for the same response) and “shuffle pairs” (Day 1 vs Day 2 of responses with same reward timing form and same conditioned interval). Box limits in box plots represent 25th and 75th percentiles, lines correspond to roughly +/-2.7σ. For demonstration purposes, outliers have been removed from plot. *** in panel C: Z = −2.73, p = 6.40 × 10^−3^ Wilcoxon rank-sum test.*** in panel D: Z = −3.61, p = 3.07 × 10^−4^.

**Figure 4, Supplemental Figure 1:**
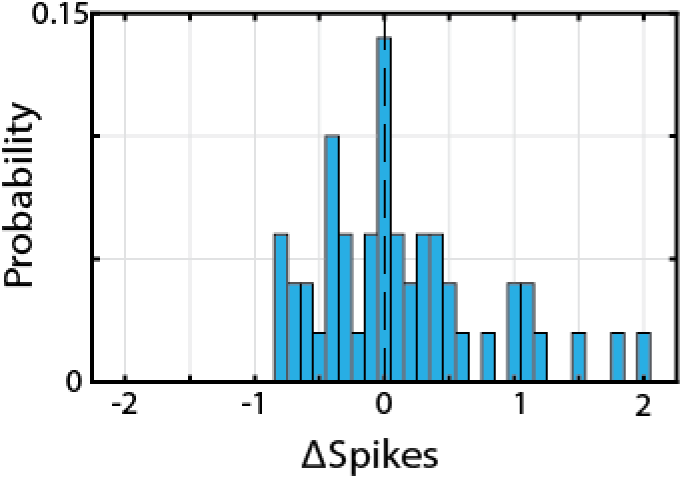
Identified interneuron activity is not influenced by licking behavior. Distribution of ΔSpikes (a variable which reflects the average change in the number of spikes before and after a first lick, see Methods) for all identified interneurons. Vertical, dashed line represents median of distributions which is not significantly different from zero (Z = 0.926, p = 0.355, Wilcoxon Signed-Rank test).

**Figure 4, Supplemental Figure 2:**
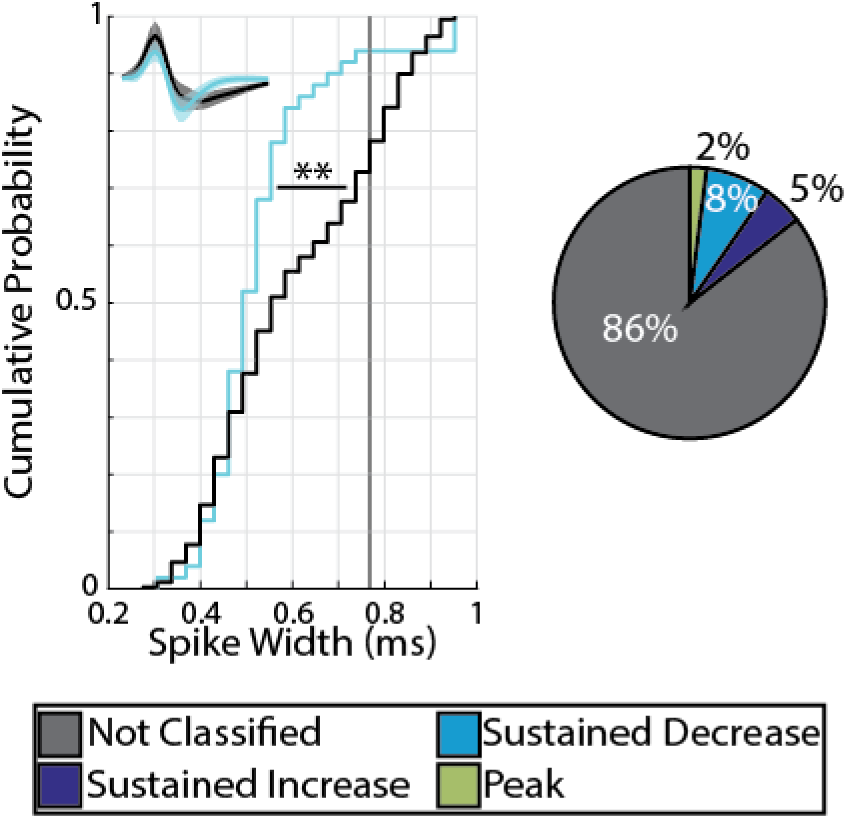
Putative pyramidal neurons express reward timing in all forms. Left: Cumulative distribution plots of calculated Spike Width for identified interneurons (cyan) and all other neurons (black). Identified interneurons have significantly narrower spike widths compared to unidentified counterparts (Z = −2.61, p = 9.2 × 10^−3^ Wilcoxon Rank-Sum test). Vertical, black line shows threshold value to define putative pyramidal cells (neurons with spike widths in the top quartile of distribution). Inset at top left shows mean± standard deviation of average waveform for identified interneurons (cyan) or putative pyramidal cells (black). Right: Pie chart showing distribution of reward timing forms within putative excitatory cells. **: p < 0.01, Wilcoxon Rank-Sum test

